# Costing ‘the’ MTD

**DOI:** 10.1101/150821

**Authors:** David C. Norris

## Abstract

**Background:** Absent adaptive, individualized dose-finding in early-phase oncology trials, subsequent registration trials risk suboptimal dosing that compromises statistical power and lowers the probability of technical success (PTS) for the investigational drug. While much methodological progress has been made toward adaptive dose-finding, and quantitative modeling of dose-response relationships, most such work continues to be organized around a concept of ‘the’ maximum tolerated dose (MTD). But a new methodology, Dose Titration Algorithm Tuning (DTAT), now holds forth the promise of *individualized* ‘MTD_*i*_’ dosing. Relative to such individualized dosing, current ‘one-size-fits-all’ dosing practices amount to a *constraint* that imposes *costs* on society. This paper estimates the magnitude of these costs.

**Methods:** Simulated dose titration as in (Norris 2017) is extended to 1000 subjects, yielding an empirical MTD_*i*_ distribution to which a gamma density is fitted. Individual-level efficacy, in terms of the probability of achieving remission, is assumed to be an E_max_-type function of *dose relative to MTD*_*i*_, scaled (arbitrarily) to identify MTD_*i*_ with the LD_50_ of the individual’s tumor. (Thus, a criterion 50% of the population achieve remission under individualized dosing in this analysis.) Current practice is modeled such that all patients receive a first-cycle dose at ‘the’ MTD, and those for whom MTD_*i*_ < MTD_the_ experience a ‘dose-limiting toxicity’ (DLT) that aborts subsequent cycles. Therapy thus terminated is assumed to confer no benefit. Individuals for whom MTD_*i*_ *≥* MTD_the_ tolerate a full treatment course, and achieve remission with probability determined by the E_max_ curve evaluated at MTD_the_/MTD_*i*_. A closed-form expression is obtained for the population remission rate, and maximized numerically over MTD_the_ as a free parameter, thus identifying the best result achievable under one-size-fits-all dosing. A sensitivity analysis is performed, using both a perturbation of the assumed Emax function, and an antipodal alternative specification.

**Results:** Simulated MTD_*i*_ follow a gamma distribution with shape parameter *α ≈* 1.75. The population remission rate under one-size-fits-all dosing at the maximizing value of MTD_the_ proves to be a function of the shape parameter—and thus the coefficient of variation (CV)—of the gamma distribution of MTD_*i*_. Within a plausible range of CV(MTD_*i*_), one-size-fits-all dosing wastes approximately half of the drug’s population-level efficacy. In the sensitivity analysis, sensitivity to the perturbation proves to be of second order. The alternative exposure-efficacy specification likewise leaves all results intact.

**Conclusions:** The CV of MTD_*i*_ determines the efficacy lost under one-size-fits-all dosing at ‘the’ MTD. Within plausible ranges for this CV, failure to individualize dosing can effectively halve a drug’s value to society. In a competitive environment dominated by regulatory hurdles, this may reduce the value of shareholders’ investment in the drug to *zero*.

**Epilogue:** The main result on one-size-fits-all dosing is generalized to regimens with several dose levels. Implications for the ongoing ALTA-1L trial are briefly explored; the 2 dose levels in the brigatinib arm of this trial may lend it a competitive advantage over the single-dose crizotinib arm.

## INTRODUCTION

Dose Titration Algorithm Tuning (DTAT), a new methodology for individualized dose-finding in early-phase oncology studies, holds forth a promise of individualized dosing from the earliest stages of oncology drug development (Norris 2017). Most immediately and obviously, such individualized dosing serves the imperative of *individual ethics* in seeking to optimize the care of each person who enrolls in a Phase I study. But by increasing the efficiency of drug development overall, individualized dosing also serves wider social aims. Less effective, ‘one-size-fits-all’ dosing may condemn valuable drugs to failure in later registration trials. More efficacious, individualized dosing may therefore avert financial losses to shareholders in pharmaceutical innovation, while preserving innovations valuable to society at large. This brief technical note estimates the magnitude of the *social* costs incurred by one-size-fits-all dose-finding studies. The argument should be of interest to shareholders in pharmaceutical innovation, and to executives having fiduciary responsibilities to them.

## THE DISTRIBUTION OF MTD_*i*_

In (Norris 2017), DTAT was demonstrated by simulated dose titration in 25 simulated subjects drawn randomly from a population model of the pharmacokinetics and myelosuppressive dynamics of docetaxel. By extending this simulation to 1000 subjects, we obtain the empirical distribution of *individualized maximum tolerated dose* (MTD_*i*_) shown in Figure 1.

**Figure 1.**
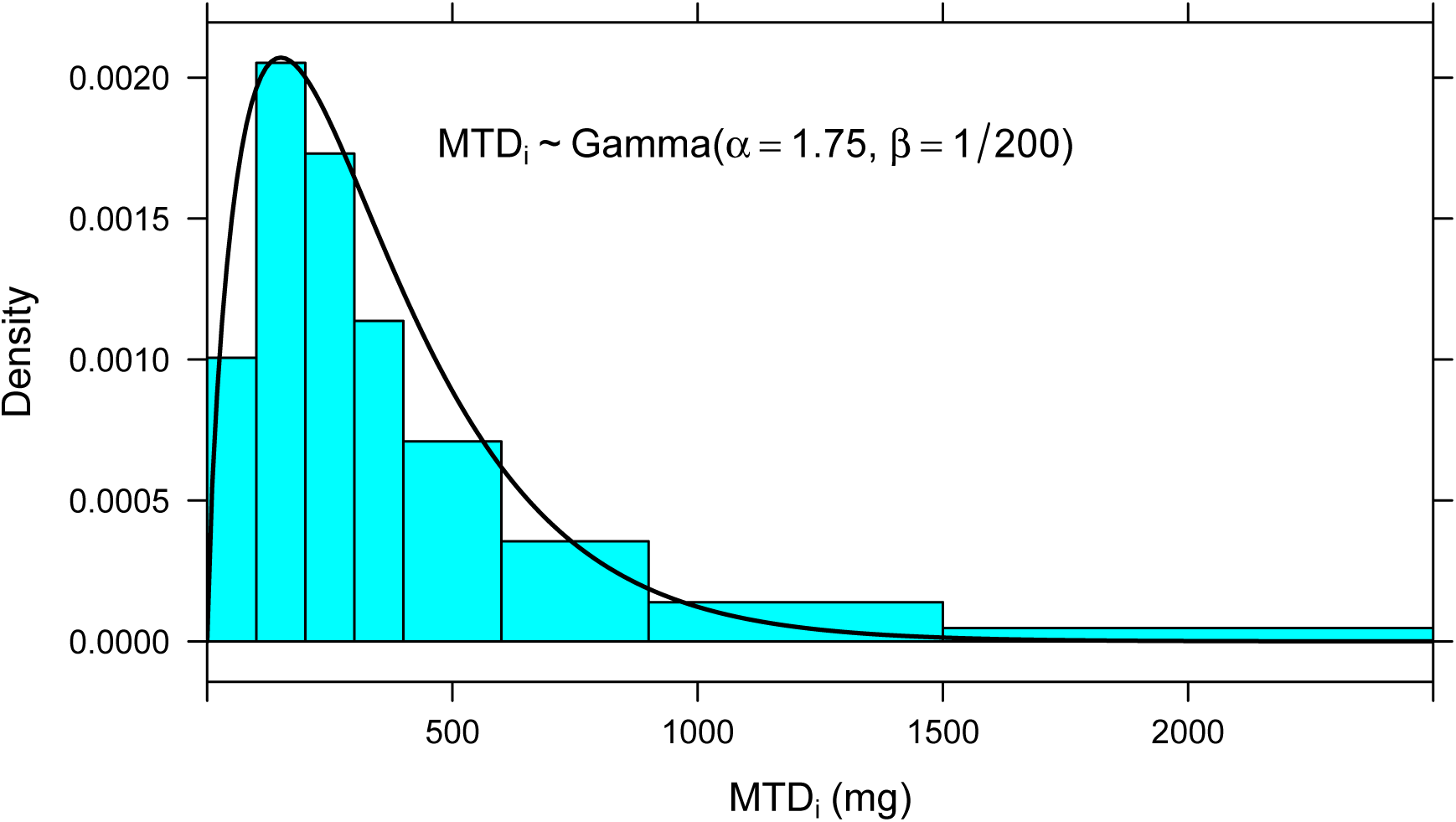
MTD_*i*_ is approximately Gamma distributed.

Whether the fitted Gamma density in Figure 1 represents a true distribution in any actual human population matters less for what follows than establishing the basic plausibility of a Gamma-distributed MTD_*i*_ generally.

## DOSE-RESPONSE MODEL

To estimate the cost of sub-MTD_*i*_ dosing, one must model individual-level efficacy as a function of dose. A traditional approach in this context is to posit a dose-effect model of a standard ‘E_max_’ type. Taking the *tumor’s* point of view, we may write in fact a ‘toxicology’ form of the model:

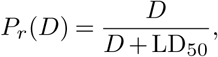

where *P*_*r*_(*D*) is the probability of achieving remission as a function of *D*, the dose received, and LD_50_ is the dose that would be ‘lethal’ to the tumor in 50% of patients—that is, the dose that would achieve remission with probability 0.5. By supposing further that MTD_*i*_ is the LD_50_ for the tumor in individual *i*, we obtain:

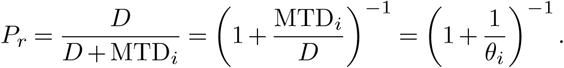

Thus, identifying MTD_*i*_ with the LD_50_ of the tumor yields a modeled remission probability that is a function of *θ*_*i*_ = *D/*MTD_*i*_, the *fraction of MTD*_*i*_ *received*. The reasonableness of this identification will be explored in the Discussion below. As it turns out, a slightly different functional form for *P*_*r*_(*θ*) supports obtaining an intermediate result in terms of standard functions:

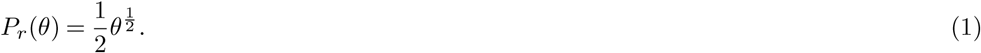

The reader suspicious of this departure from tradition should take reassurance in noting that this revised functional form is uniformly *more forgiving* of suboptimal dosing than the standard form:

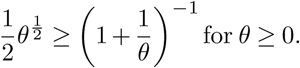

## THE DISTRIBUTION OF *θ*_*i*_ = MTD_the_*/*MTD_*i*_

If MTD_*i*_ *∼* Gamma(*α, β*), then 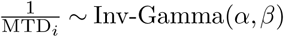 and consequently

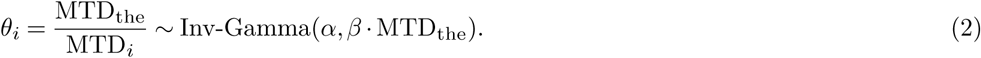

## THE TWO COSTS OF ONE-SIZE-FITS-ALL DOSING

Under the prevailing practice of one-size-fits-all dosing at ‘the’ MTD, we take the following to occur: (1) Those individuals *i* for whom MTD_*i*_ *>* MTD_the_ will receive suboptimal dosing at a fraction *θ*_*i*_ *<* 1 of their optimal dose; (2) those for whom MTD_*i*_ *<* MTD_the_ will experience intolerable adverse effects with a first dose, and will not receive subsequent cycles of therapy. (Those rare individuals for whom MTD_*i*_ = MTD_the_ holds *exactly* will receive their optimal *θ*_*i*_ = 1 dose, and enjoy the full benefit of the drug.) Thus, dosing everyone at MTD_the_ has two social costs: individuals who cannot tolerate ‘the’ MTD derive no benefit from the drug, while those who could have tolerated higher doses derive suboptimal benefit. This latter cost is well described in a literature stretching back 2 decades, documenting (across many types of cancer) that patients who experience milder adverse effects from chemotherapy tend to have worse outcomes (Saarto et al. 1997, Cameron et al. (2003), Di Maio et al. (2005), Yamanaka et al. (2007), Y. H. Kim et al. (2009), Lee et al. (2011), Shitara et al. (2011), McTiernan et al. (2012), Liu, Zhang, and Li (2013), Shiozawa et al. (2014), Su et al. (2015), Osorio et al. (2017)).

Figure 2 depicts the balance of these costs under 3 different choices of MTD_the_ *∈* {100, 200, 300} mg. If MTD_*i*_ *∼* Gamma(*α* = 1.75*, β* = 1*/*200), then *θ*_*i*_ = MTD_the_*/*MTD_*i*_ will follow the inverse gamma distribution:

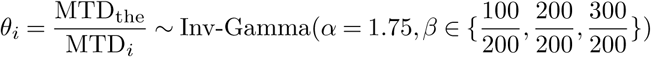

These 3 densities are plotted in green in Figure 2, superimposed on the dose-response relationship of Equation 1. Here, it is readily seen that setting MTD_the_ = 100 mg causes most individuals to receive doses below half of their MTD_*i*_’s (*θ <* 0.5). Conversely, setting MTD_the_ = 300 mg causes few individuals to be dosed at *θ ≤* 0.5, but excludes a large fraction of the population from treatment—as indicated by the large area under the dashed curve to the right of *θ* = 1.

**Figure 2.**
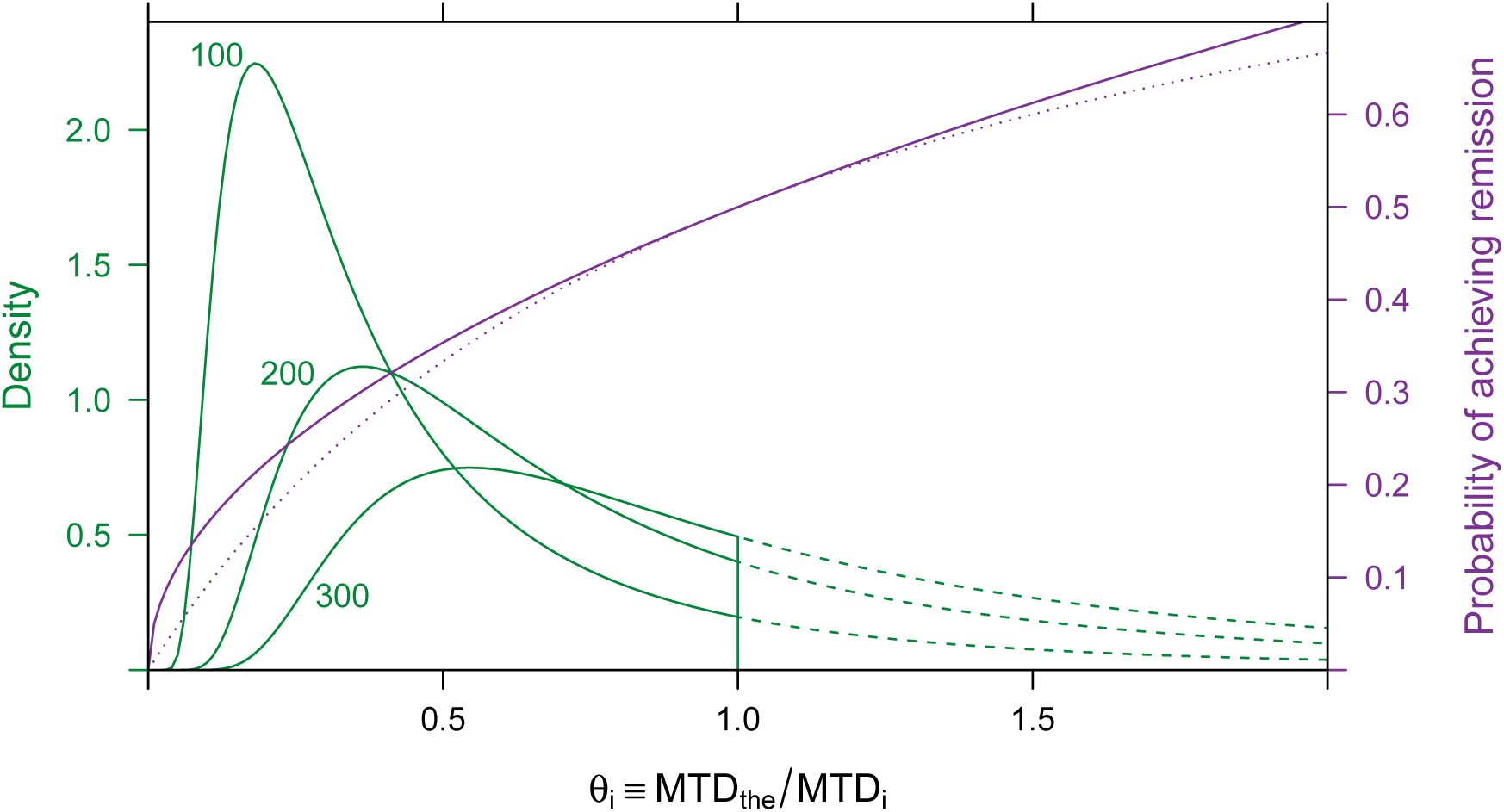
Social costs of one-size-fits-all dosing at 3 different choices of ‘the’ MTD. Against the purple dose-response function, the distribution of *θ*_*i*_ = MTD_the_*/*MTD_*i*_ is plotted for 3 different values of MTD_the_. When ‘the’ MTD is set low (100 mg), few individuals are excluded from treatment (area under dashed curves), but most are treated at a low fraction (*θ*_*i*_ *<* 0.5) of their MTD_*i*_’s. Conversely, when ‘the’ MTD is set high (300 mg), fewer individuals are dosed so low, but many (large area under dashed curve) cannot tolerate the drug and do not receive a full course of treatment.

## POPULATION-LEVEL EFFICACY OF ONE-SIZE-FITS-ALL DOSING

Given that *θ*_*i*_ is distributed as in Equation 2, and that the individual-level probability of remission is as given by Equation 1, then the *population rate* 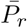 of achieving remission may be calculated by integrating *P*_*r*_(*θ*_*i*_) over the treated population 0 *≤ θ*_*i*_ *≤*1. Normalizing 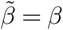. MTD_the_, we can calculate:

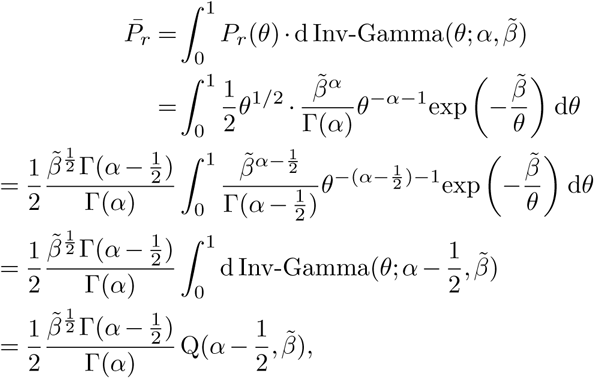

where *Q* denotes the regularized gamma function.

The best-case population rate of remission is obtained by choosing MTD_the_ optimally:

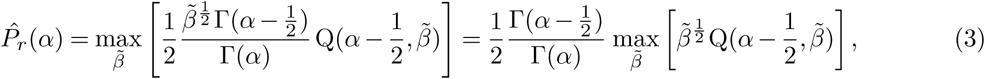

in which it should be noted particularly that 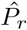 is a function of the ‘shape parameter’ *α*, which determines the *coefficient of variation* (CV) of our gamma-distributed MTD_*i*_ via CV = *α*^*-*1*/*2^. The maximand on the right-hand side of Equation 3 is readily evaluated using the implementation of the regularized gamma function *Q* provided in R package **zipfR** (Evert and Baroni 2007), and the maximum obtained numerically. The dependence of 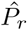 on CV(MTD_*i*_) is plotted in Figure 3.

**Figure 3.**
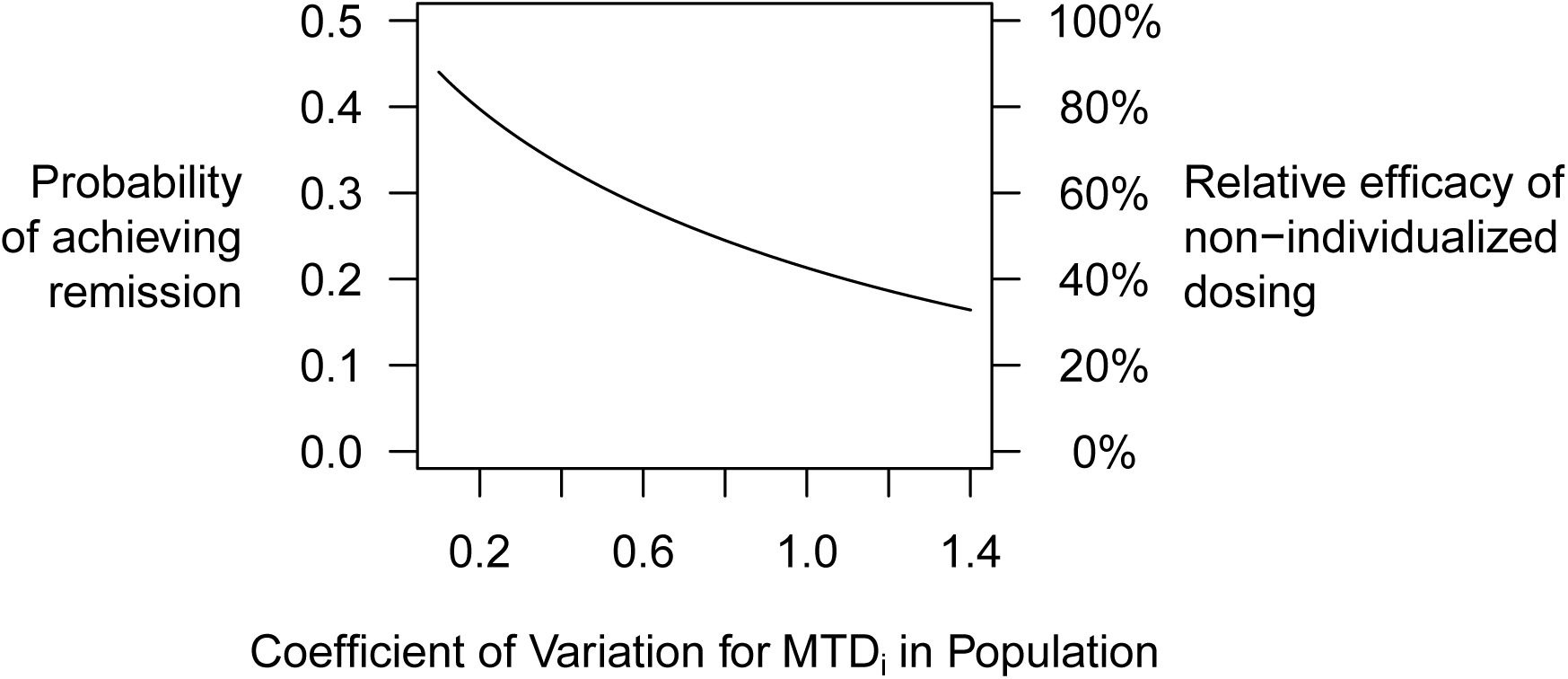
Estimated *minimum* cost of one-size-fits-all dosing, as a function of the coefficient of variation (CV) of MTD_*i*_ in the population. It is assumed that ‘the’ MTD is chosen to maximize the population-level remission rate, under the constraint of one-size-fits-all dosing. The cost of the one-size-fits-all constraint is calculated relative to a reference remission probability of 50% for optimal individualized dosing at each patient’s MTD_*i*_. The more MTD_*i*_ varies within the population, the more untenable one-size-fits-all dosing becomes.

## SENSITIVITY ANALYSIS

One assumption essential to the development of my argument thus far was that individual-level outcomes are a function of *θ*_*i*_ = *D/*MTD_*i*_, the fraction of MTD_*i*_ received by individual *i*. This assumption would hold in the limiting case where inter-individual variation in MTD_*i*_ was driven entirely by *pharmacokinetic* heterogeneity. (Consider the particularly simple example of an oral drug for which otherwise-identical individuals differed only regarding *bioavailability*.) Nevertheless, the sensitivity of my results to this assumption does seem to warrant further examination.

Consider the following perturbation of Equation 1:

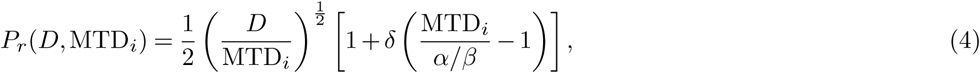

where the factor in brackets induces a dependence of *P*_*r*_ on MTD_*i*_ that is not accounted for by *θ*_*i*_ = *D/*MTD_*i*_. This factor is motivated as a first-order Taylor expansion of a general functional dependence, centered on the population mean E[MTD_*i*_] = *α/β*. Not only do we recover Equation 1 in the limit as *δ →* 0, but we also preserve *independently of δ* the same population-average remission rate of 1*/*2 under individualized ‘MTD_*i*_’ dosing. (To appreciate this latter point, set *D* = MTD_*i*_ in Equation 4 then take expectations on both sides.)

To obtain 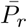 as previously, we rearrange Equation 4 as follows

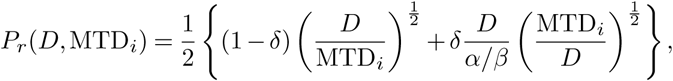

and then integrate as before:

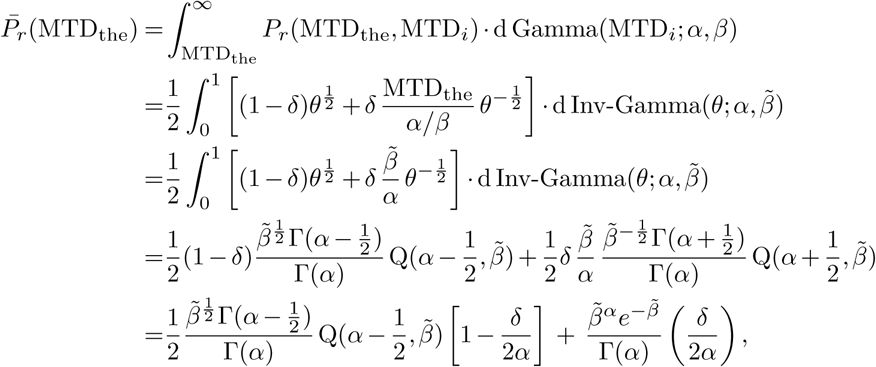

in which the final step requires applications of the recurrence relations for Γ and Q.

The best-case population rate of remission is obtained, as before, by choosing MTD_the_ optimally:

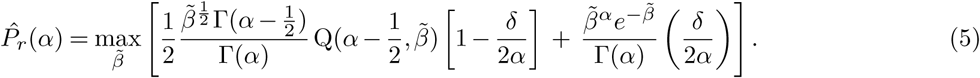

Equation 5 generalizes Equation 3, and reduces to it on setting *δ* = 0.

Without attempting to develop an interpretation of *δ*, we plot in Figure 4 the relative efficacy of one-size-fits-all dosing at selected values of *CV*, for *δ ∈* [–1, 1]. Evidently, the perturbation is of second order. Moreover, absolute deviations from the picture of Figure 3 prove small over the range of what seems from Equation 4 to be the natural scale for *δ*.

**Figure 4.**
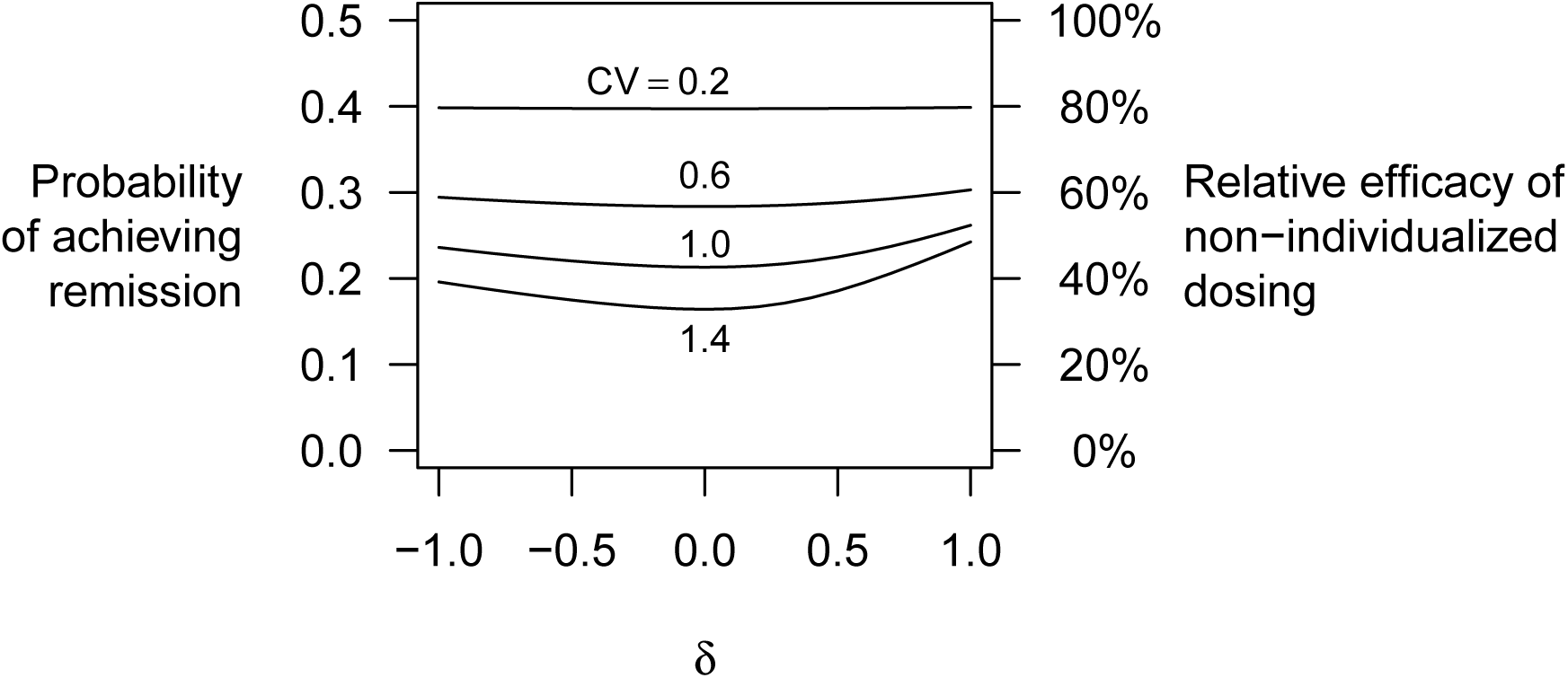
Sensitivity of efficiency loss estimate to perturbation as in Equation 4.

**Figure 5.**
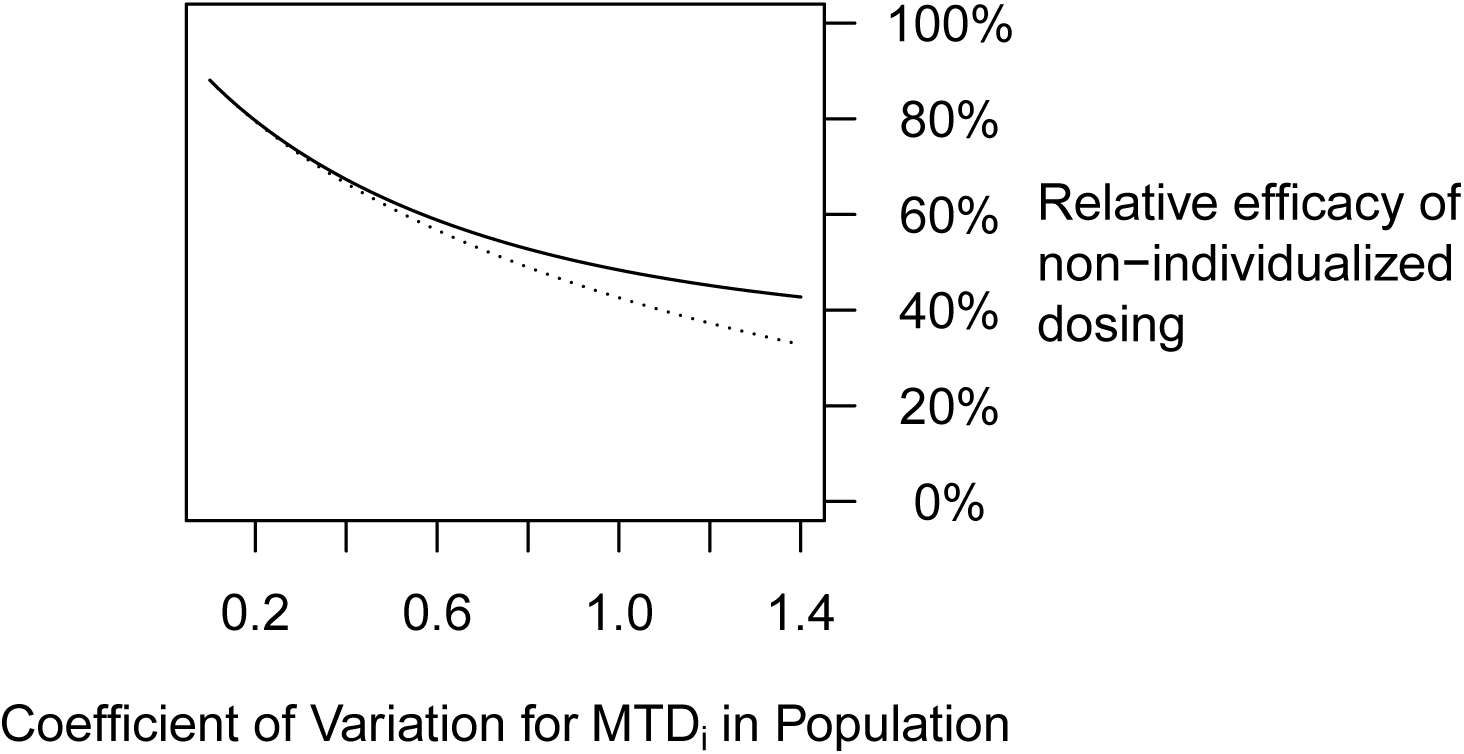
Sensitivity of Figure 3 to alternative specification of an MTD_*i*_-independent dose-effect relation as in Equation 6. The solid curve shows efficacy loss under this alternative specification, while the dotted curve shows, for comparison, the original curve derived from Equation 1. Evidently, this alternative specification does little to redeem one-size-fits-all dosing.

### Sensitivity under an interpretable alternative

As an alternative to the foregoing perturbation analysis, we might instead posit a single, readily interpretable alternative to Equation 1. A possibility that immediately presents itself as a ‘diametrically opposed’ alternative is to suppose *P*_*r*_(*D,* MTD_*i*_) *independent of* MTD_*i*_, as in:

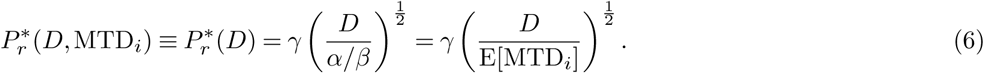

Under one-size-fits-all dosing, Equation 6 yields a population-level efficacy of

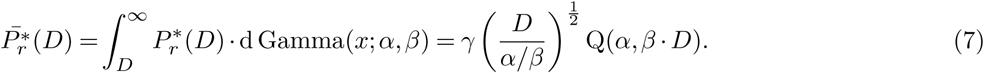

In contrast to the cases considered above, here the integration extends into a region where *P*_*r*_ *>* 1; but, by taking *γ →* 0, we can push this region as far as desired into the tail of our Gamma-distributed MTD_*i*_. (As will be seen presently, *γ* drops out of our analysis of *relative* efficacy.) Differentiating with respect to *D*, and employing the same normalization 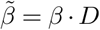 introduced previously, we find that Equation 7 is maximized at 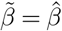 defined by

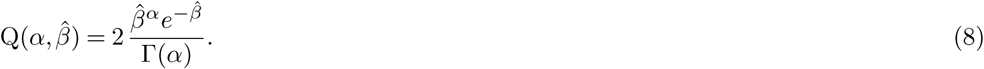

Substituting (8) into (7), we find that this maximum value is:

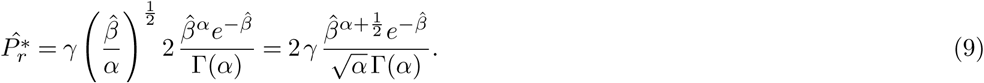

Under the MTD_*i*_-independent remission probability (6), optimal individualized dosing does not reduce trivially to a constant 1*/*2. Rather, we must calculate as follows:

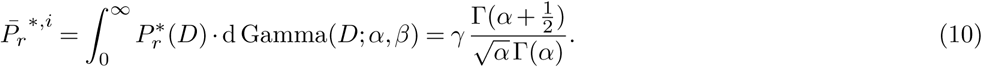

Dividing (9) by (10), we obtain the following expression for relative efficacy of one-size-fits-all dosing under Equation 6:

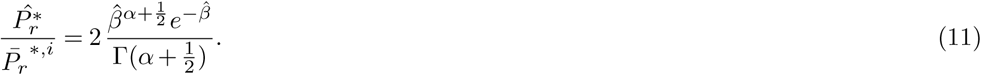

Solving Equation 8 numerically for 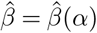, and substituting this into Equation 11, we obtain the following counterpart to Figure 3:

## DISCUSSION

At two points in this argument, I have adopted modeling assumptions that are relatively *forgiving* of one-size-fits-all dosing, and therefore would tend to underestimate its costs. Firstly, my highly concave square-root E_max_ model (Equation 1) regards under-dosing more favorably than does a typical E_max_ model, such as appears in the dotted purple curve in Figure 2. Secondly, the optimization itself in Equation 3 surely overestimates the population-level outcomes achieved by one-size-fits-all dosing as implemented in current Phase I designs. Indeed, these designs tend to target DLT *rates* without explicit reference or regard to outcomes.

Dose reduction protocols, as seen both in trials and in clinical practice, do somewhat relax the extreme form of one-size-fits-all dosing constraint that I have modeled in this paper. Clearly, such protocols exist precisely to recoup some part of the lost value I calculate here. But given that these protocols are readily interpreted as a (very) poor man’s DTAT, their existence only underscores the urgent need for rational dose individualization in oncology. This conclusion fully withstands the rather vigorous sensitivity analysis performed above.

## CONCLUSIONS

Taking *population-level efficacy* as a proxy, I have estimated the social cost of one-size-fits-all dosing organized around a concept of ‘the’ maximum tolerated dose (MTD) in oncology. The magnitude of this cost is seen to depend primarily on the coefficient of variation (CV) of *individually optimal* MTD_*i*_ doses in the population. Within plausible ranges for this CV, the failure to individualize dosing can effectively halve a drug’s value to society. Notably, in a competitive environment dominated by regulatory hurdles, this may reduce the value of shareholders’ investment in a drug to *zero*.

## DATA AVAILABILITY

*Open Science Framework:* Data for Figure 1 may be found in R package **DTAT** (v0.1-1), available together with code for reproducing all of this paper’s Figures and analyses, at doi: 10.17605/osf.io/vtxwq.

### Grant information

The author declared that no grants were involved in supporting this work.

## EPILOGUE

The reader who has followed my argument up to this point will have no difficulty generalizing Equation 3 to the case of optimal titration confined to *n* fixed doses 0 ≡ *D*_0_ *< D*_1_ *< … < D*_*n*_:

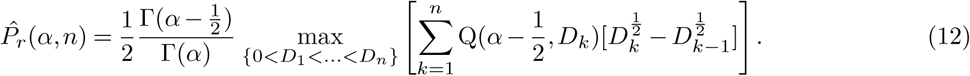

In fact, the case *n* = 2 immediately appertains to a Phase III trial of current interest, ALTA-1L, which incorporates a two-dose titration procedure in one arm (Ariad Pharmaceuticals 2017). In this trial comparing brigatinib and crizotinib in ALK-positive NSCLC, patients in the brigatinib arm will receive the drug at 90 mg daily for 7 days, with subsequent escalation (as tolerated) to 180 mg daily. Such a protocol immediately raises the question as to how much of the ‘wasted value’ documented above may be recoverable by a simple protocol that escalates through a small number of equally-spaced doses. For the sake of simplicity and definiteness, let us replace the *n*-dimensional simplex over which Equation (12) maximizes by the 1-dimensional space of dose levels defined by 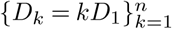, regarding *D*_1_ as the sole free parameter:

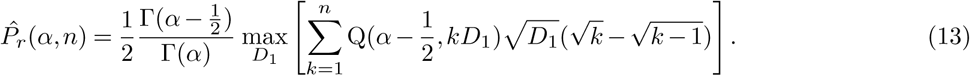

This yields Figure 6, generalizing Figure 3 to include the cases of *n* = 2, 3, or 6 dose levels. Apparently, even modest departures from one-size-fits-all dosing may recoup substantial amounts of efficacy. Over a middling range of CV in Figure 6, adding just 2 dose levels (for a total of 3) boosts relative efficacy by 20%. It seems conceivable that the dose titration employed in the brigatinib arm of ALTA-1L lends this arm a competitive advantage over the single-dose crizotinib arm.

**Figure 6.**
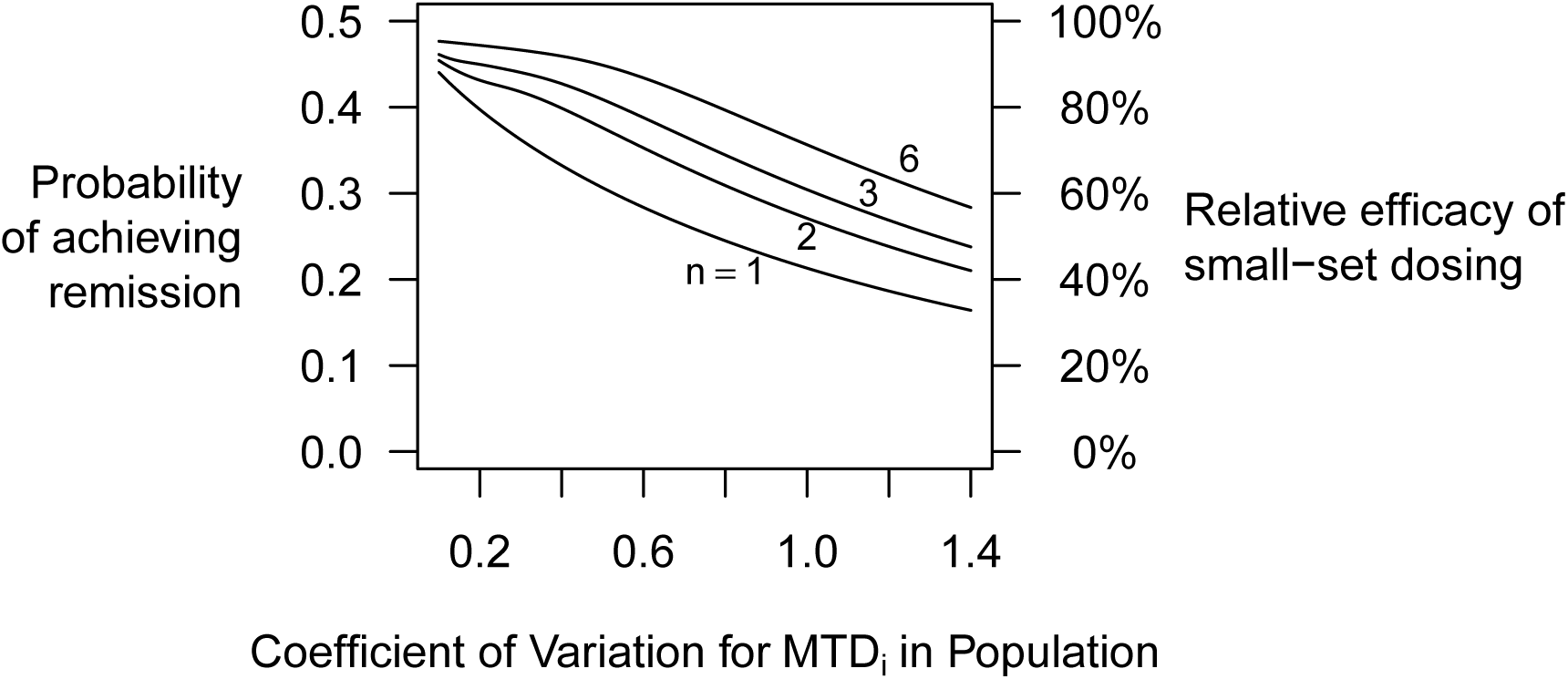
Partial recovery of the value lost to one-size-fits-all dosing by expanding the number of available doses to *n* = 2, 3 and 6 levels. For simplicity, we consider dose sets constructed from a basic dose *D*_1_ and its integer multiples up to *D*_*n*_ = *nD*_1_. For the sake of definiteness, the free parameter *D*_1_ of each such dose set is chosen—conditional on *n* and CV(MTD_*i*_)—to maximize the population-level remission rate achievable with that set. In the case of drugs with moderate inter-individual variation in MTD_*i*_, even a handful of (well chosen) dose levels suffices to achieve reasonably efficient dose individualization.

## REFERENCES

Ariad Pharmaceuticals. 2017. “ALTA-1l: A Phase 3 Study of Brigatinib Versus Crizotinib in ALK-Positive Advanced Non-Small Cell Lung Cancer Patients [Last Updated: 2017 Aug 10].” In. Clinical-Trials.gov [Internet]. Bethesda, MD: U.S. National Library of Medicine. https://clinicaltrials.gov/ct2/show/NCT02737501.

Cameron, D. A., C. Massie, G. Kerr, and R. C. F. Leonard. 2003. “Moderate Neutropenia with Adjuvant CMF Confers Improved Survival in Early Breast Cancer.” British Journal of Cancer 89 (10): 1837–42. doi:10.1038/sj.bjc.6601366.

Di Maio, Massimo, Cesare Gridelli, Ciro Gallo, Frances Shepherd, Franco Vito Piantedosi, Silvio Cigolari, Luigi Manzione, et al. 2005. “Chemotherapy-Induced Neutropenia and Treatment E°cacy in Advanced Non-Small-Cell Lung Cancer: A Pooled Analysis of Three Randomised Trials.” The Lancet. Oncology 6 (9): 669–77. doi:10.1016/S1470-2045(05)70255-2.

Evert, Stefan, and Marco Baroni. 2007. “*zipfR*: Word Frequency Distributions in R.” In Proceedings of the 45th Annual Meeting of the Association for Computational Linguistics, Posters and Demonstrations Sessions, 29–32. Prague, Czech Republic.

Kim, Yun Hwan, Hyun Hoon Chung, Jae Weon Kim, Noh-Hyun Park, Yong-Sang Song, and Soon-Beom Kang. 2009. “Prognostic Significance of Neutropenia During Adjuvant Concurrent Chemoradiotherapy in Early Cervical Cancer.” Journal of Gynecologic Oncology 20 (3): 146–50. doi:10.3802/jgo.2009.20.3.14>6.

Lee, C. K., H. Gurney, C. Brown, R. Sorio, N. Donadello, G. Tulunay, W. Meier, et al. 2011. “CarboplatinPaclitaxel-Induced Leukopenia and Neuropathy Predict Progression-Free Survival in Recurrent Ovarian Cancer.” British Journal of Cancer105 (3): 360–65. doi:10.1038/bjc.2011.256.

Liu, Wei, Cui-Cui Zhang, and Kai Li. 2013. “Prognostic Value of Chemotherapy-Induced Leukopenia in Small-Cell Lung Cancer.” Cancer Biology& Medicine10 (2): 92–98. doi:10.7497/j.issn.2095-3941.2013.02.005.

McTiernan, Anne, Rachel C. Jinks, Matthew R. Sydes, Barbara Uscinska, Jane M. Hook, Martine van Glabbeke, Vivien Bramwell, et al. 2012. “Presence of Chemotherapy-Induced Toxicity Predicts Improved Survival in Patients with Localised Extremity Osteosarcoma Treated with Doxorubicin and Cisplatin: A Report from the European Osteosarcoma Intergroup.” European Journal of Cancer (Oxford, England: 1990)48 (5): 703–12. doi:10.1016/j.ejca.2011.09.012.

Norris, David C. 2017. “Dose Titration Algorithm Tuning (DTAT) Should Supersede 'the’ Maximum Tolerated Dose (MTD) in Oncology Dose-Finding Trials.” F1000Research 6 (July): 112. doi:10.12688/f1000research.10624.3.

Osorio, J. C., A. Ni, J. E. Chaft, R. Pollina, M. K. Kasler, D. Stephens, C. Rodriguez, et al. 2017. “Antibody-Mediated Thyroid Dysfunction During T-Cell Checkpoint Blockade in Patients with Non-Small-Cell Lung Cancer.” Annals of Oncology. Accessed March 6. doi:10.1093/annonc/mdw640.

Saarto, T., C. Blomqvist, P. Rissanen, A. Auvinen, and I. Elomaa. 1997. “Haematological Toxicity: A Marker of Adjuvant Chemotherapy E°cacy in Stage II and III Breast Cancer.” British Journal of Cancer75 (2): 301–5.

Shiozawa, Yusuke, Junko Takita, Motohiro Kato, Manabu Sotomatsu, Katsuyoshi Koh, Kohmei Ida, and Yasuhide Hayashi. 2014. “Prognostic Significance of Leukopenia in Childhood Acute Lymphoblastic Leukemia.” Oncology Letters7 (4): 1169–74. doi:10.3892/ol.2014.1822.

Shitara, Kohei, Keitaro Matsuo, Isao Oze, Ayako Mizota, Chihiro Kondo, Motoo Nomura, Tomoya Yokota, Daisuke Takahari, Takashi Ura, and Kei Muro. 2011. “Meta-Analysis of Neutropenia or Leukopenia as a Prognostic Factor in Patients with Malignant Disease Undergoing Chemotherapy.” Cancer Chemotherapy and Pharmacology68 (2): 301–7. doi:10.1007/s00280-010-1487-6.

Su, Zhen, Yan-Ping Mao, Pu-Yun OuYang, Jie Tang, Xiao-Wen Lan, and Fang-Yun Xie. 2015. “Leucopenia and Treatment E°cacy in Advanced Nasopharyngeal Carcinoma.” BMC Cancer 15 (May): 429. doi:10.1186/s12885-015-1442-3.

Yamanaka, T., S. Matsumoto, S. Teramukai, R. Ishiwata, Y. Nagai, and M. Fukushima. 2007. “Predictive Value of Chemotherapy-Induced Neutropenia for the E°cacy of Oral Fluoropyrimidine S-1 in Advanced Gastric Carcinoma.” British Journal of Cancer97 (1): 37–42. doi:10.1038/sj.bjc.6603831.

